# Which life history strategy can maintain high genetic diversity in plants?

**DOI:** 10.1101/2022.04.26.489613

**Authors:** Yoichi Tsuzuki, Takenori Takada, Masashi Ohara

## Abstract

Classifying a variety of plant life histories is an important step to understand ecological and evolutionary background of life history diversification in plants. Principal component analysis (PCA) of life history traits, as well as elasticity analysis using a ternary diagram, successfully categorized the diverse life histories and clarified their link to various ecological characteristics, such as population dynamics, functional traits, and conservation status. However, the relationships with population genetic properties, including genetic diversity, have not been explored very much. In this study, we overlaid annual change rate of expected heterozygosity *η* on life history spectrum obtained by PCA and elasticity analysis to examine which life history strategy can maintain high genetic diversity over time. We found that *η* gradually changed along the fast-slow continuum and that slow-paced life histories maintained high genetic diversity, probably due to generation overlap. Elasticity analysis showed that life histories that highly depend on stasis also maintained genetic diversity. As genetic diversity reflects adaptive potential to environmental changes, our study provides a new genetic perspective on adaptative evolution and population viability to life history study.

## Background

Most plant life history traits, such as longevity (annual vs biennial vs perennial) and degree of iteroparity (semelparous vs iteroparous), are different among populations and species [1, 2]. This variation translates into a high diversity of plant life history, or lifetime schedule of survival, growth, and reproduction. The array of life histories and their population dynamics can be modelled with matrix population models (MPMs), which divide life cycle into several stage classes and describe lifetime trajectory from birth to death using transition probabilities and fecundity of each stage class [3].

Plant MPMs have been compared among populations and species to understand ecological and evolutionary background of life history diversification. One approach is correlation analysis of multiple life history traits obtained from MPMs [4, 5, 6]. Because life history traits are correlated with one another due to trade-off among survival, growth, and reproduction [7], ordination such as principal component analysis (PCA) can summarize multiple life history traits to a couple of dimensions. Two major axes turned out to explain life history variation across taxa worldwide: the fast-slow continuum and the reproductive strategy [4, 6]. Meanwhile, elasticity analysis, in which elasticities of population growth rate (*λ*) to stage-dependent transition probabilities and fecundities have been calculated from MPMs, has also been used to classify life history strategies. Elasticity of *λ* to a matrix element *a_ij_* (*ij*-th element that denotes the flow of individuals from *j*-th to *i*-th stages) is defined as the relative change in *λ* compared to the relative change in *a_ij_* and represents the relative contribution of *a_ij_* to the population growth rate *λ*.

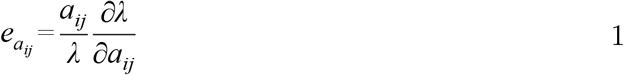

Because the sum of elasticities over all matrix elements is equal to unity [8], life history can be plotted on a ternary diagram by taking partial sum of elasticities for three demographic processes: fecundity, growth, stasis (defined as survival without progressive growth) (Figure 1a) [9, 10, 11, 12]. It was shown that the position on the ternary diagram reflects life form (Figure 1b-d) [9]. These classifications enabled us to overlay a variety of variables, including population growth rate, functional traits, environmental conditions, and conservation status, onto life history spectrum [6, 11, 12, 13], facilitating evaluation on life history strategies from various ecological, evolutionary, and conservation points of view.

**Figure 1.**
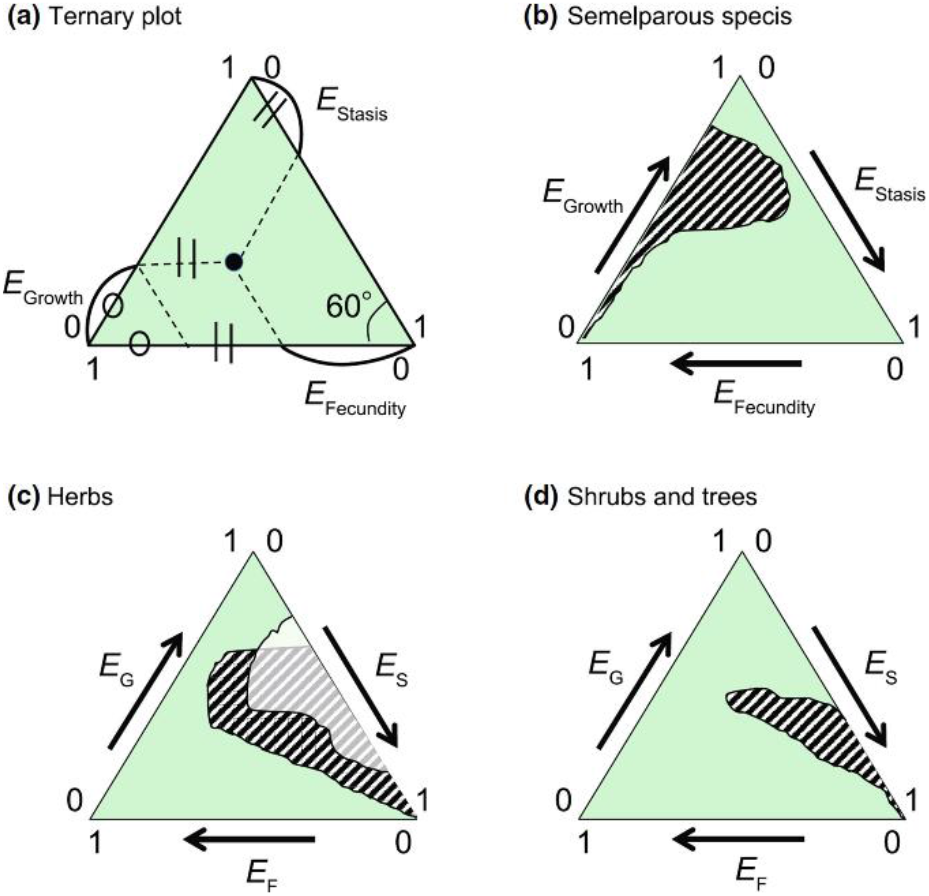
A Ternary diagram of the elasticity for 84 wild plant species’ populations. The three axes correspond to the sum of the elasticities of three main demographic processes: fecundity (F), growth (G) and stasis (S). (a) Coordinates in the ternary plot; (b) area occupied by semelparous species, (c) area occupied by herbs in open habitats (hatched) and in forest habitats (white), and (d) area occupied by shrubs and trees. Source: Figure 1 of [11] (Reproduced with permission of Wiley).

Meanwhile, life history is known to influence genetic properties of a population, such as genetic diversity [14, 15, 16, 17], fixation probability of a mutation [18, 19], and the speed of adaptation [20, 21]. By linking these genetic properties to life history spectrum, we could figure out the overall relationships between life history strategies and their genetic consequences. Especially, genetic diversity reflects the potential to adapt to future environmental changes and is often used as an important proxy of population viability [22, 23], being an important index to be assessed across a variety of life history strategies. Although there have been several attempts [15, 16, 24], previous studies handled only a few life history characteristics, such as longevity and breeding system. The whole relationships between the complex life history spectrum and genetic diversity remained unclear.

In this study, we employed a theoretical approach to figure out which life history strategies are likely to maintain genetic diversity in plants. We made two-stage life histories in silico, covering wide range of life history strategies. We estimated annual change rate of expected heterozygosity *η*, which was developed in [25], and examined its relationships with a life history spectrum obtained by PCA and elasticity analysis.

## Methods

### Model

We considered a life history model with two stages, juvenile (stage 1) and adult (stage 2), supposing a perennial plant species (Figure 2). We defined four transition probabilities (*t*_11_, *t*_21_, *t*_12_, and *t*_22_) and fecundity (*f*_12_) per year. These five stage-dependent vital rates were classified into three groups following [10] and [11]: growth (*t*_21_), stasis (*t*_11_, *t*_12_, *t*_22_), and fecundity (*f*_12_). Matrix population model can be formulated as follows:

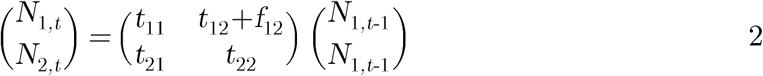

where *N*_1,*t*_ and *N*_2,*t*_ are the number of individuals in stage 1 and 2 in year *t*, respectively. The dominant eigenvalue of the population projection matrix (i.e., the 2 × 2 matrix in the right-hand side) is population growth rate *λ* [3].

**Figure 2.**
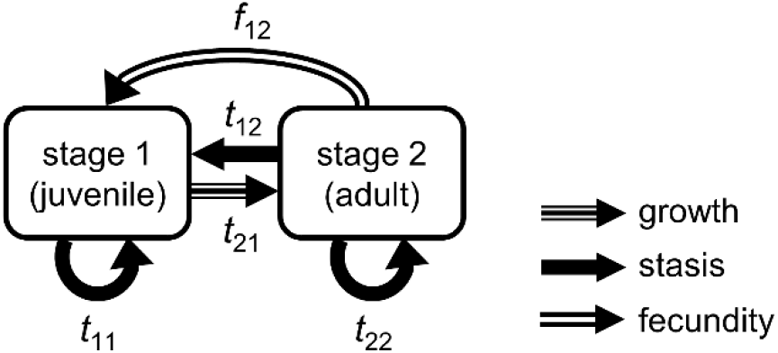
The two-stage life history used in this study. The probability of growth (*t*_21_), stasis (*t*_11_ and *t*_22_), and retrogression (*t*_12_) determines the transition of individuals per year. Annual fecundity is denoted as *f*_12_. The line types of the arrows represent the classification of [10, 11].

### Annual change rate of expected heterozygosity (*η*)

[25] theoretically derived *η*, which is the annual change rate of expected heterozygosity in stage-structured plant populations at an equilibrium state, where population size and stage distribution remain constant overtime. Expected heterozygosity is a common proxy of genetic diversity, defined as the probability that two genes randomly sampled from a population is non-identical-by-descent, which means that the two genes do not have a common ancestor. In [25], the differential equation of expected heterozygosity in a two-stage life history was formulated as follows:

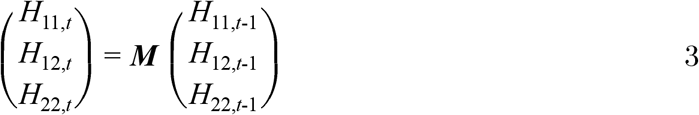

where *H_ij,t_* is the probability that two genes, each sampled from stage *i* and *j* in year *t*, are non-identical-by-descent. The dominant eigenvalue of the 3 × 3 matrix ***M*** in the right-hand side, which is denoted as *η* hereafter, is annual change rate of expected heterozygosity, in the same manner as *λ* being population growth rate in Equation (2). Nine elements of ***M*** are formulated with stage-dependent vital rates (*t*_21_, *t*_11_, *t*_12_, *t*_22_, and *f*_12_) and the number of individuals in each stage (*N*_1,*t*_ and *N*_2,*t*_), which are shown in detail in the supplementary material 1.

The four transition probabilities were set to be either of 0, 0.19, 0.38, 0.57, 0.76, and 0.95, with three constraints: (1) 0 ≤ *t*_11_ + *t*_21_ ≤ 1 and 0 ≤ *t*_12_ + *t*_22_ ≤ 1 (sum of probabilities should range from 0 to 1), (2) *t*_21_ ≠ 0 (if *t*_21_ = 0, the flow of individuals from stage 1 to 2 is stopped and the life cycle is broken off), (3) *t*_11_ + *t*_22_ > 0 (at least either of the stasis probabilities is greater than zero, otherwise genes in different stages are completely isolated, leading to one population eternally divided into two subpopulations). Assuming the equilibrium state, we adjusted *f*_12_ to satisfy *λ* = 1, *N*_1,*t*_ = *N*_1,*t*-1_, and *N*_2,*t*_ = *N*_2,*t*-1_. Total population size *N* was set to 100, and we split *N* into stage 1 and 2 as proportional to stable stage distribution, that is, the leading right eigenvector of the population projection matrix in Equation (2). A total of 285 combinations of parameter values were generated, for each of which we obtained *η*.

### Life history traits

We calculated eight life history traits: generation time (*T*), survivorship curve type (*H*), age at sexual maturity (*L_α_*), mean probability of progressive growth (*γ*) and retrogressive growth (*ρ*), the degree of iteroparity (*S*), mean reproduction rate (*φ*), and mature life expectancy (*L_ω_*). Each life history trait is related to either of the four demographic processes: *T* is related to turnover, while *H* and *L_α_* to survival, *γ* and *ρ* to growth, and *S, φ,* and *L_ω_* to reproduction, respectively. These traits were estimated following the method of [3] and [6]. In estimating *T, H*, and *S*, we followed [26] to transform stage-dependent demographic rates to age-dependent ones. The detailed calculation procedures are explained in supplementary material 2. Generation time *T* was estimated as the expected age of parents of a newborn cohort (Equation S24). Survivorship curve type *H* is Keyfitz’ entropy (Equation S26), which is a continuous variable that reflects the shape of survivorship curve from type I (high mortality at mature ages, *H* < 1), II (constant mortality throughout the lifespan, *H* = 1), to III (high mortality at young ages, *H* > 1). Age at sexual maturity *L_α_* is the average age to become sexually reproductive (Equation S19). Mean probability of progressive and retrogressive growth (*γ* and *ρ*) and mean reproduction rate (*φ*) are the number of individuals that grow, retrogress, and are born per year divided by the total population size, respectively (Equation S28-S30). The degree of iteroparity *S* represents the temporal spread of reproductive events along lifespan and is quantified by Demetrius’ entropy (Equation S25). Mature life expectancy *L_ω_* is estimated by subtracting *L_α_* from life expectancy of a newborn (Equation S20). We carried out principal component analysis (PCA) of the eight life history traits and examined how the resultant principal components are correlated with *η*.

### Elasticity analysis

For the 200 parameter combinations, we obtained elasticity of *λ* to the five stage-dependent vital rates, using a function “elasticity” in an R package “popbio” [27]. The function “elasticity” does not return elasticities of *t*_12_ and *f*_12_, while it outputs that of *a*_12_ (= *t*_12_ + *f*_12_). We used the following equations to obtain elasticities of *t*_12_ and *f*_12_.

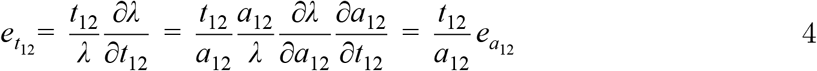

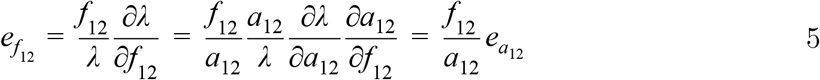

Equation (4) and (5) shows that the elasticity of *a*_12_ is proportionally allocated to *t*_12_ and *f*_12_, assuring that the basic property of elasticity that the overall sum is equal to unity holds after obtaining *e*_*t*_12__ and *e*_*f*_12__. We took a partial sum of the five elasticities by their categories: growth, stasis, and fecundity.

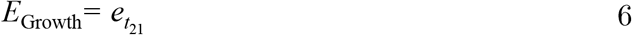

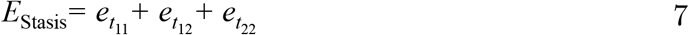

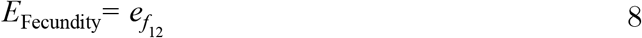

We drew a ternary plot whose three axes were *E*_Growth_, *E*_stasis_, and *E*_Fecundity_. We examined how the positions on the ternary diagram are related to *η*.

All analyses were carried out in R version 4.1.1 [28]. The R code is available at supplementary material 4.

## Results

### Dependence of *η* on transition probabilities

*η* increased with the four transition probabilities and was especially large under maximum *t*_11_ and *t*_22_ (Figure 3a). By tracking small squares along the axes of *t*_21_ and *t*_12_, we could see that *η* became gradually high with increasing *t*_21_ and *t*_12_, although the increment was subtle compared to the increase along the two stasis probabilities.

**Figure 3.**
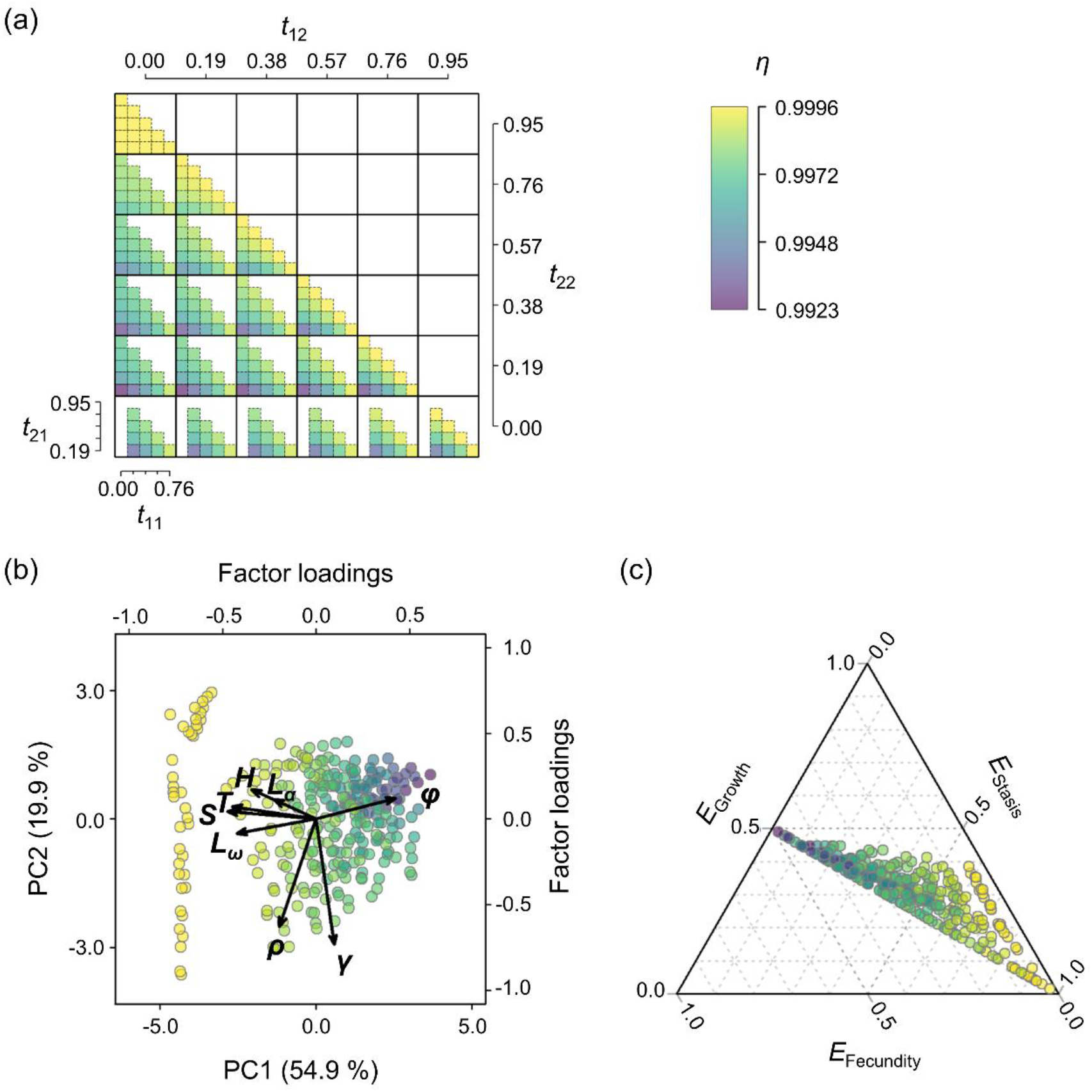
(a) Parameter dependence of *η* on the four transition probabilities is shown in a nested manner. Rows and columns of large square correspond to *t*_22_ and *t*_12_, respectively. The positions of small squares within a large one represent *t*_21_ and *t*_11_. The scale of the axes is given with the interval of 0.19. (b) PCA plot of the eight life history traits: generation time (*T*), survivorship curve type (*H*), age at sexual maturity (*L_α_*), mean probability of progressive growth (*γ*) and retrogressive growth (*ρ*), the degree of iteroparity (*S*), mean reproduction rate (*φ*), and mature life expectancy (*L_ω_*). (c) Ternary diagram of elasticity partially summed into fecundity (*E*_Fecundity_), growth (*E*_Growth_), and stasis (*E*_Stasis_). For all the three panels, the colour filled in squares or dots represent the value of *η*.

### Life history traits and *η*

PCA yielded two axes that could jointly explain 74.8 % of the total variation in the eight life history traits (Figure 3b, S1). Judging from the factor loadings, the first axis PC1 was positively correlated with mean reproduction rate *φ* and negatively with generation time *T*, degree of iteroparity *S*, mature life expectancy *L_ω_*, age at sexual maturity *L_α_*, and survivorship curve type *H*. In other words, PC1 reflected the gradient from long-lived (large *T* and *L_ω_*), slow-paced (large *L_α_*), and iteroparous (large *S*) life histories with low fecundity (low *φ*) and high mortality risk during early ages (large *H*), to short-lived, fast-paced, highly reproductive, and semelparous ones with high mortality at mature ages. This correlation exactly corresponds to “the fast-slow continuum,” which describes that high mortality risk after sexual maturation is related to fast growth, high fecundity, and short lifespan [5, 29]. The second axis PC2 was negatively correlated with progressive and retrogressive growth (*ρ* and *γ*).

*η* increased along the gradient from fast-to slow-paced life history (from large to small PC1 scores, Figure 3b). In other words, *η* was large under low annual fecundity, long generation time, delayed sexual maturation, high degree of iteroparity, long life expectancy after maturation, and survivorship curve type 3 (concentrated death at the beginning of the life cycle). PC2, which was constituted by progressive and retrogressive growth, did not show strong correlation with *η*: *η* neither increased nor decreased apparently along the vertical direction of the PCA plot (Figure 3b).

### Elasticity and *η*

The 285 life histories we made in silico were distributed in the center and the lower-right region of the ternary plot (Figure 3c), which overlapped with the distribution of herbs, shrubs, and trees (Figure 1c, 1d). The distribution region corresponded to 0 ≤ *E*_Fecundity_ ≤ 0.5, 0 ≤ *E*_Growth_ ≤ 0.5, and 0 ≤ *E*_Stasis_ ≤ 1. Overall, *η* gradually increased with decreasing *E*_Fecundity_ (i.e., increased in the horizontal direction from left to right), and was especially large in life histories that occupied the right edge of the triangle, where *E*_Stasis_ was relatively high (0.5 < *E*_Stasis_ < 1) compared to *E*_Fecundity_ and *E*_Growth_.

## Discussion

### Fast-slow continuum and *η*

For the 285 life histories we made in silico, PCA yielded two major axes that explained the variation of the eight life history traits. The first axis (PC1) reflected the fast-slow continuum, which was composed of *T, H, L_α_, S, φ*, and *L_ω_*. Contrary to our results, recent analyses using empirical MPMs showed that progressive growth *γ* also constituted the fast-slow continuum while the degree of iteroparity *S* formed another independent axis regarding reproductive strategy [4, 6]. In this study, we only handled the equilibrium state where population size and stage distribution are constant over time to meet the assumption of *η*, while empirical MPMs contained non-equilibrium populations. The assumption of constant population size balances mortality and recruitment and causes trait collinearity [30], which could have been one reason of different life history traits aligned with the fast-slow continuum. Despite the discrepancy, the fact that the major axis explaining life history variation is the fast-slow continuum holds in our analysis.

We found that *η* increased with decreasing PC1 scores (Figure 3b), indicating that high *η* is realized in slow-paced life histories. The reason why slow-paced strategies maintain higher genetic diversity can be attributed to generation overlap. Generation overlap enables individuals with different age to temporally coexist, which facilitates the accumulation of genetic variation [31]. The prominent positive effect of stasis probabilities (*t*_11_ and *t*_22_, Figure 3a) would also reflect the contribution of generation overlap, because stasis would keep old individuals to remain in the same stage, delaying generation turnover. Although the contribution of longevity or stasis to genetic diversity had been suggested [14, 15, 17, 24], most studies handled only a limited number of species. Our results successfully supported their generality in the overall life history spectrum.

The second axis (PC2) was comprised of two life history traits related to growth. Considering that PC2 did not have apparent correlation with *η*, it can be said that both progressive and retrogressive growth were almost neutral to the gradient of *η*. This result indicates that inter-stage transitions do not affect the dynamics of genetic diversity very much. This is in line with the result that growth and retrogression probabilities (*t*_21_ and *t*_12_) had less contribution to *η* than stasis probabilities (Figure 3a).

### Elasticity and *η*

Although elasticity has been used to clarify key demographic processes for population growth, its relationship with genetic dynamics remained unclear. Our study successfully bridged the gap and found that *η* was high in the bottom-right area of the ternary plot, where large proportion of elasticity was assigned to stasis (Figure 3c). This region is occupied by iteroparous herbs and woody plants (Figure 1c, 1d) [9] and is characterized by long lifespan [32]. Thus, this result indicates that long lifespan is related to high *η*, which is in line with the result of PCA.

Because *E*_stasis_ reflects the relative impact of perturbation in stasis probabilities (*t*_11_, *t*_12_, and *t*_22_) on population growth rate [8], we could say that population dynamics of life histories with high *η* is strongly determined by stasis probabilities while not so much by growth probability (*t*_21_) and fecundity (*f*_12_). While growth and reproduction can jointly accelerate generation turnover and replacement of gene pool, stasis promotes survival of established individuals and the maintenance of preexisting genes. High dependency of population dynamics on stasis probably should have contributed to the maintenance of existing genetic variation, leading to high *η*.

### Assessing population viability from genetic point of view

Both PCA and elasticity analysis clarified that slow-paced life history with heavy dependency on stasis can maintain genetic diversity over the course of time. Genetic diversity is a prerequisite for adaptation to novel environments [33, 34] and plays a crucial role in population persistence under environmental changes [22]. Thus, our study enables us to examine viable life history strategies in terms of evolutionary potential, suggesting *η* as a promising index of population viability along with conventionally used *λ* (population growth rate).

Still, there remains problems to be solved. In this study, we took a theoretical approach that did not involve empirical data. COMPADRE and COMADRE, which are the databases of empirically estimated MPMs [35, 36], have been used to classify various plant and animal life history strategies [4, 6, 37]. Matrices available at these databases include non-equilibrium ones, assuming exponential population growth or decline. Because the number of individuals determines the strength of genetic drift, or stochastic loss of genetic variation [38], *η* covaries with changing population size time by time under a non-equilibrium state, hardly being a good predictor of long-term genetic dynamics. Thus, the annual change rate of expected heterozygosity *η* and its relationship with life history spectrum would be mostly applicable to a short time scale. To solve the incompatibility with preexisting empirical life history data and to realize long-term forecast, an improved modelling or statistics which can handle varying *N*, as well as varying *η*, should be developed.

## Funding

This research was financially supported by Japan Society for the Promotion of Science KAKENHI [grant no. 19H03294, 20K06821, and 21J10814].

## Notes

### Competing Interest Statement

The authors have declared no competing interest.

