## Supplemental Material for "Which life history strategy can maintain high genetic diversity in plants?"

Corresponding author: Yoichi Tsuzuki

N10W5, Kita-ku, Sapporo City, Hokkaido Prefecture, Japan, 060-0810

phone number: (+81)-011-706-2225

#### Contents

**1 Detailed elements of matrix *M*** **2**

**2 Calculating life history traits** **4**

2.1 Age at sexual maturity ($L_\alpha$) and mature life expectancy ($L_\omega$) . . . . . . . . . . . 4

2.2 Generation time ($T$), the degree of iteroparity ($S$), and survivorship curve type ($H$) 5

2.3 Mean probability of progressive and retrogressive growth ($\gamma$ and $\rho$) and mean reproduction rate ($\phi$) . . . . . . . . . . . . . . . . . . . . . . . . . . . . . 6

**3 Explanatory power of principal components of the eight life history traits** **8**

**4 R code** **9**

### 1 Detailed elements of matrix $\boldsymbol{M}$

Here, we show the elements of matrix $\boldsymbol{M}$ in Equation (3) of the main text under the equilibrium state. Because we consider an equilibrium state where population size and stage distribution remain constant over the course of time, we denote $N_{1,t}$ and $N_{2,t}$ as $N_1$ and $N_2$, respectively. The equations below are quoted from Tsuzuki et al. (2021).

$$\boldsymbol{H_t} = \begin{pmatrix} H_{11,t} \\ H_{12,t} \\ H_{22,t} \end{pmatrix} = \boldsymbol{M} \begin{pmatrix} H_{11,t-1} \\ H_{12,t-1} \\ H_{22,t-1} \end{pmatrix}$$

$$= \begin{pmatrix} m_{11} & m_{12} & m_{13} \\ m_{21} & m_{22} & m_{23} \\ m_{31} & m_{32} & m_{33} \end{pmatrix} \begin{pmatrix} H_{11,t-1} \\ H_{12,t-1} \\ H_{22,t-1} \end{pmatrix} \tag{S1}$$

where

$$m_{11} = t_{11}^2 \frac{1 - 1/(2t_{11}N_1)}{1 - 1/(2N_1)} \tag{S2}$$

$$m_{21} = \frac{N_1}{N_2} \left( \frac{t_{11}t_{21}}{1 - 1/(2N_1)} \right) \tag{S3}$$

$$m_{31} = \left( \frac{t_{21}N_1}{N_2} \right)^2 \frac{1 - 1/(2t_{21}N_1)}{1 - 1/(2N_1)} \tag{S4}$$

$$m_{12} = \frac{2t_{11}a_{12}N_2}{N_1} \tag{S5}$$

$$m_{22} = t_{11}t_{22} + a_{12}t_{21} \tag{S6}$$

$$m_{32} = \frac{2t_{21}t_{22}N_1}{N_2} \tag{S7}$$

$$m_{13} = \left(\frac{t_{12}N_2}{N_1}\right)^2 \frac{1-1/(2t_{12}N_2)}{1-1/(2N_2)} + \frac{2t_{12}f_{12}N_2^2}{N_1^2} + \left(\frac{f_{12}N_2}{N_1}\right)^2 \left(1-\frac{1}{2f_{12}N_2}\right) \tag{S8}$$

$$m_{23} = \frac{N_2}{N_1}\left(\frac{t_{12}t_{22}}{1-1/(2N_2)} + f_{12}t_{22}\right) \tag{S9}$$

$$m_{33} = t_{22}^2 \frac{1-1/(2t_{22}N_2)}{1-1/(2N_2)} \tag{S10}$$

#### 2 Calculating life history traits

##### 2.1 Age at sexual maturity ($L_\alpha$) and mature life expectancy ($L_\omega$)

Firstly, we decomposed the population projection matrix into two: $\boldsymbol{U}$ matrix, which describes the transition process, and $\boldsymbol{F}$ matrix, which is made up of stage-specific fecundity:

$$\begin{pmatrix} t_{11} & t_{12} + f_{12} \\ t_{21} & t_{22} \end{pmatrix} = \begin{pmatrix} t_{11} & t_{12} \\ t_{21} & t_{22} \end{pmatrix} + \begin{pmatrix} 0 & f_{12} \\ 0 & 0 \end{pmatrix} = \boldsymbol{U} + \boldsymbol{F} \tag{S11}$$

We calculated the fundamental matrix $\boldsymbol{N}$ following Equation (5.5) to (5.7) in Caswell (2001).

$$\boldsymbol{N} = (\boldsymbol{E} - \boldsymbol{U})^{-1} \tag{S12}$$

$$= \begin{pmatrix} n_{11} & n_{12} \\ n_{21} & n_{22} \end{pmatrix} \tag{S13}$$

where $\boldsymbol{E}$ is the identity matrix. Fundamental matrix provides the time to absorption state, that is, the state from which individuals will not move. In the case of Equation (S12) and (S13), absorption state is death. Life expectancy of a newborn $L$, or the time to death, is equal to the sum of the first column of $\boldsymbol{N}$.

$$L = n_{11} + n_{21} \tag{S14}$$

Next, we added stage 2 as another absorption state, and obtained corresponding fundamental matrix $\boldsymbol{N'}$.

$$\boldsymbol{N'} = \left( \begin{pmatrix} 1 & 0 \\ 0 & 1 \end{pmatrix} - \begin{pmatrix} t_{11} & 0 \\ t_{21} & 0 \end{pmatrix} \right)^{-1} \tag{S15}$$

$$= \begin{pmatrix} n'_{11} & n'_{12} \\ n'_{21} & n'_{22} \end{pmatrix} \tag{S16}$$

We then calculated conditional fundamental matrix $\boldsymbol{N}^{(\boldsymbol{C})}$, with condition on absorption in stage 2, following Equation (5.24) of Caswell (2001).

$$\boldsymbol{N}^{(\boldsymbol{C})} = \begin{pmatrix} n'_{21} & 0 \\ 0 & n'_{22} \end{pmatrix}^{-1} \begin{pmatrix} n'_{11} & n'_{12} \\ n'_{21} & n'_{22} \end{pmatrix} \begin{pmatrix} n'_{21} & 0 \\ 0 & n'_{22} \end{pmatrix} \tag{S17}$$

$$= \begin{pmatrix} n_{11}^{(C)} & n_{12}^{(C)} \\ n_{21}^{(C)} & n_{22}^{(C)} \end{pmatrix} \tag{S18}$$

We can obtain age at sexual maturity $L_\alpha$ as the sum of the first column of $\boldsymbol{N}^{(\boldsymbol{C})}$.

$$L_\alpha = n_{11}^{(C)} + n_{21}^{(C)} \tag{S19}$$

Subsequently, mature life expectancy can be obtained as follows:

$$L_\omega = L - L_\alpha \tag{S20}$$

#### 2.2 Generation time ($T$), the degree of iteroparity ($S$), and survivorship curve type ($H$)

By multiplying $\boldsymbol{U}$ matrix $x$ times, we could obtain transition probabilities per $x$ years.

$$\boldsymbol{U}^x = \begin{pmatrix} t_{11} & t_{12} \\ t_{21} & t_{22} \end{pmatrix}^x = \begin{pmatrix} \tilde{u}_{11} & \tilde{u}_{12} \\ \tilde{u}_{21} & \tilde{u}_{22} \end{pmatrix} \tag{S21}$$

Here, $\tilde{u}_{11}$ and $\tilde{u}_{21}$ are the probabilities that an individual in stage 1 remain in stage 1, or move to stage 2, after $x$ years, respectively. Now, we can formulate age-specific survival rate $l_x$, which denotes the probability of a newborn individual to survive until age $x$, and age-specific fecundity $m_x$, which is a expected number of newborns that an individual of age $x$ can make.

$$l_x = \tilde{u}_{11} + \tilde{u}_{21} \tag{S22}$$

$$m_x = 0 \times \frac{\tilde{u}_{11}}{\tilde{u}_{11} + \tilde{u}_{21}} + t_{21} \times \frac{\tilde{u}_{21}}{\tilde{u}_{11} + \tilde{u}_{21}} \tag{S23}$$

Then, we obtained generation time ($T$), the degree of iteroparity ($S$), and survivorship curve type ($H$), using the following equations.

$$T = \frac{\sum_{x=1}^{x_{max}} x l_x m_x}{\sum_{x=1}^{x_{max}} l_x m_x} \tag{S24}$$

$$S = -\sum_{x=1}^{x_{max}} e^{-log\lambda} l_x m_x log(e^{-log\lambda} l_x m_x) \tag{S25}$$

$$H = \frac{\sum_{x=1}^{x_{max}} log(l_x) l_x}{\sum_{x=1}^{x_{max}} l_x} \tag{S26}$$

where $x_{max}$ is the maximum age defined as the age at which either of the two criteria (quoted from Waples et al., (2013)) is satisfied.

1. oldest age for which $l_x$ was $\geq 1$ % of the value at age at maturity ($L_\alpha$)

2. oldest age for which the product $l_x v_x$ was $\geq 1$ % of the maximum $l_x v_x$ for any age, where $v_x$ is the reproductive value of an individual of age $x$

Equation (S24) was quoted from Equation (5.75) in Caswell (2001), while Equation (S25) and (S26) were from Table S2 in the extended methods and materials of Salguero-Gómez et al. (2016).

##### 2.3 Mean probability of progressive and retrogressive growth ($\gamma$ and $\rho$) and mean reproduction rate ($\phi$)

We obtained stable stage distribution by calculating the leading right eigenvector (denoted as $\boldsymbol{w}$ hereafter) of the population projection matrix.

$$\boldsymbol{w} = \begin{pmatrix} w_1 \\ w_2 \end{pmatrix} \tag{S27}$$

$\gamma$, $\rho$, and $\phi$ were defined by the following equations:

$$\gamma = \frac{t_{21} w_1}{w_1 + w_2} \tag{S28}$$

$$\rho = \frac{t_{12} w_2}{w_1 + w_2} \tag{S29}$$

$$\phi = \frac{f_{12} w_2}{w_1 + w_2} \tag{S30}$$

#### 3 Explanatory power of principal components of the eight life history traits

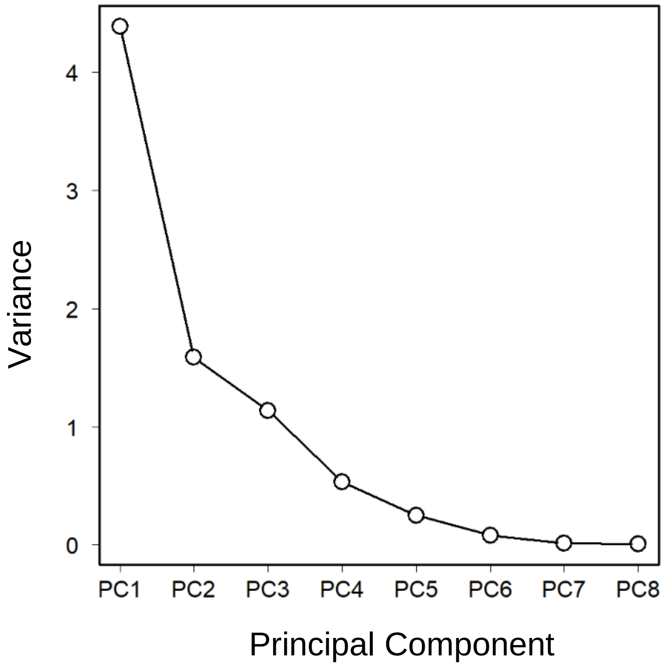


**Figure S1** Variance explained by each principal component of the eight life history traits

### 4 R code

```
####################################################################
# Which life history strategy can maintain high genetic diversity?
#
#   R Script written by Yoichi Tsuzuki
#
####################################################################

##-------------------------------------
## 1. Create life histories
##-------------------------------------

library(gtools)

parameter_combinations <-
  matrix(rep(c(1,-1),21),2)*
  t(combinations(6,2,0:5*0.19,repeats.allowed = T))+
  matrix(rep(c(0,0.95),21),2)

demrates0 <- cbind(apply(parameter_combinations,1,rep,each=dim(parameter_combinations)[2]),
                   apply(parameter_combinations,1,rep,times=dim(parameter_combinations)[2]))
demrates0 <- as.data.frame(demrates0)
names(demrates0) <- c("t11","t21","t12","t22")

demrates <- subset(demrates0,
                   t11+t21 <= 1 & t12+t22 <= 1 & # probabilities <= 1
                     t21 > 0 &
                     t11 + t22 > 0)

demrates$f12 <- (1-demrates$t11)*(1-demrates$t22)/demrates$t21-demrates$t12
demrates$a12 <- demrates$t12+demrates$f12

summary(demrates);dim(demrates)

##-------------------------------------
## 2. Obtain eta
##-------------------------------------

# 1. stable stage structure

Nsize <- 100

Nstructure <- NULL
lambda <- NULL
for(i in 1:dim(demrates)[1]){
  matrix.i <- matrix(c(demrates$t11[i],demrates$t21[i],demrates$a12[i],demrates$t22[i]),2)
  structure.i <- abs(eigen(matrix.i)$vectors[,eigen(matrix.i)$values==max(eigen(matrix.i)$values)])
  Nstructure <- rbind(Nstructure,structure.i/sum(structure.i)*Nsize)
  lambda <- c(lambda, max(eigen(matrix.i)$values))
}
summary(lambda) # confirm that all lambda (population growth rate) is equal to 1
summary(Nstructure)

# 2. eta

eta <- NULL
for(j in 1:dim(demrates)[1]){
  H.mat.j <- matrix(c(demrates$t11[j]*(demrates$t11[j]-(1-demrates$t11[j])/(2*Nstructure[j,1]-1)),
                      (demrates$t11[j]*demrates$t21[j]*Nstructure[j,1])/
                        (Nstructure[j,2])/(1-1/(2*Nstructure[j,1])),
                      (demrates$t21[j]*(Nstructure[j,1]/Nstructure[j,2])^2)*
                        (demrates$t21[j]-(1-demrates$t21[j])/(2*Nstructure[j,1]-1)),
                      2*demrates$t11[j]*demrates$a12[j]*Nstructure[j,2]/Nstructure[j,1],
                      (demrates$t11[j]*demrates$t22[j]+demrates$t21[j]*demrates$a12[j]),
                      2*demrates$t21[j]*demrates$t22[j]*Nstructure[j,1]/Nstructure[j,2],
                      (demrates$t12[j]*(Nstructure[j,2]/Nstructure[j,1])^2)*
                        (demrates$t12[j]-(1-demrates$t12[j])/(2*Nstructure[j,2]-1))+
                        (demrates$f12[j]*(Nstructure[j,2]/Nstructure[j,1])^2)*
                        (demrates$f12[j]-1/(2*Nstructure[j,2]))+
                        2*demrates$f12[j]*demrates$t12[j]*(Nstructure[j,2]/Nstructure[j,1])^2,
                      (demrates$t12[j]*demrates$t22[j]/(1-1/(2*Nstructure[j,2]))+
                         demrates$f12[j]*demrates$t22[j])*Nstructure[j,2]/Nstructure[j,1],
                      demrates$t22[j]*(demrates$t22[j]-(1-demrates$t22[j])/(2*Nstructure[j,2]-1))),3,3)
  eigen.j <- eigen(H.mat.j)
  a.i <- (apply(Im(eigen.j$vectors),2,sum)==0) * (abs(Re(eigen.j$vectors)[1,])>0) *
    (abs(Re(eigen.j$vectors)[3,])>0) ==1
  eta.i <- max(Re(eigen.j$values[a.i]))
  eta <- c(eta,eta.i)
}

summary(eta);length(eta);hist(eta)


# 3. parameter dependency

library(tagcloud)
library(viridis)

cols.eta <- smoothPalette(eta,(viridis(1000,alpha=.6)))

plot_position <- cbind(rep(0:5*0.19,times=6),rep(0:5*0.19,each=6))

par(mar=c(0,0,0,0),oma=c(3,3,4,4))
plot(NA,xlim=c(0,max(demrates$t12)/0.19+1),ylim=c(0,max(demrates$t22)/0.19+1),
     bty="n",xaxt="n",yaxt="n",xlab="",ylab="")
```

```
for(i in 1:dim(plot_position)[1]){
  plot_position_i <- which((round(demrates$t22,2)==round(plot_position[i,1],2)) *
                           (round(demrates$t12,2)==round(plot_position[i,2],2)) == 1)
  rect(plot_position[i,2]/0.19*1+demrates$t11[plot_position_i]*0.2/0.19,
       plot_position[i,1]/0.19*1+demrates$t21[plot_position_i]*0.2/0.19-0.2,
       plot_position[i,2]/0.19*1+demrates$t11[plot_position_i]*0.2/0.19+0.2,
       plot_position[i,1]/0.19*1+demrates$t21[plot_position_i]*0.2/0.19,
       col=cols.eta[plot_position_i],border=T,lty=2)
}
segments(rep(0,6),0:6,rep(6,6),0:6,lwd=3)
segments(0:6,rep(0,6),0:6,rep(6,6),lwd=3)
axis(1,1:5/5-0.1,labels=F, lwd=2)
axis(2,1:5/5-0.1,labels=F, lwd=2)
axis(3,1:6-0.5,labels=F, lwd=2)
axis(4,1:6-0.5,labels=F, lwd=2)
par(mar=c(5.1,4.1,4.1,2.1),oma=c(0,0,0,0))

# legend
plot(NA,NA,xlim=c(0,1),ylim=c(0,1),xaxt="n",yaxt="n",xlab="",ylab="")
rect(rep(0.45,100),0.2+c(0:99)*0.6/100,rep(0.55,100),0.2+c(1:100)*0.6/100,
     col=(viridis(100,alpha=0.6)),border=(viridis(100,alpha=0.005)))
rect(0.45,0.2,0.55,0.8,col=NULL,border=1,lwd=1)
segments(0.58,0.2,0.58,0.8,lwd=3)
segments(0.58,seq(0.2,0.8,length=4),0.6,seq(0.2,0.8,length=4),lwd=3)
round(seq(min(eta),max(eta),length=4),4)
range(eta)


##-----------------------------------------
## 3. The number of individuals in each age class
##-----------------------------------------

library(expm)

N_t <- NULL # the number of individuals aged t
pb <- txtProgressBar(min=1,max=dim(demrates)[1],style=3)
for(i in 1:dim(demrates)[1]){
  matrix.i <- matrix(c(demrates$t11[i],demrates$t21[i],demrates$t12[i],demrates$t22[i]),2)
  N_t.i <- NULL
  for (y in 1:1000){
    N_t.i <- c(N_t.i,sum(matrix.i%^%(y-1) %*% matrix(c(0,0,demrates$f12[i],0),2) %*% matrix(Nstructure[i,],2)))
  }
  N_t <- rbind(N_t, N_t.i)
  setTxtProgressBar(pb,i)
}

##------------------------------------
## 4. Calculation of life history traits
##------------------------------------

# 1. age-specific survival rate

lx <- t(apply(N_t,1,function(x) x/x[1]))

# 2. calculate life expectancy (lifee), La, and Lw

lifee <- NULL; La <- NULL; Lw <- NULL
pb <- txtProgressBar(min=1,max=dim(demrates)[1],style=3)
for(i in 1:dim(demrates)[1]){
  matrix.i <- matrix(c(demrates$t11[i],demrates$t21[i],demrates$t12[i],demrates$t22[i]),2,2)
  fund.i <- solve(diag(1,2)-matrix.i)
  lifee.i <- sum(fund.i[,1])
  F.i <- matrix(c(0,0,demrates$f12[i],0),2,2)
  B.i <- matrix(c(1-sum(matrix.i[,1]),0,0,1),2,2) %*% solve(diag(1,2)-cbind(matrix.i[,1],0))
  La.i <- sum(solve(diag(1,2)-(solve(diag(B.i[2,])) %*% cbind(matrix.i[,1],0) %*% diag(B.i[2,])))[,1])
  Lw.i <- lifee.i-La.i
  lifee <- c(lifee,lifee.i)
  La <- c(La,La.i)
  Lw <- c(Lw,Lw.i)
  setTxtProgressBar(pb,i)
}

# 3. calculate gamma, rho, & phi

gamma <- NULL; phi <- NULL; rho <- NULL
for(i in 1:dim(demrates)[1]){
  matrix.i <- matrix(c(demrates$t11[i],demrates$t21[i],
                       demrates$a12[i],demrates$t22[i]),2,2)
  strc.i <- abs(eigen(matrix.i)$vectors[,eigen(matrix.i)$values==max(eigen(matrix.i)$values)])
  gamma.i <- strc.i[1]/sum(strc.i)*demrates$t21[i]
  phi.i <- strc.i[2]/sum(strc.i)*demrates$f12[i]
  rho.i <- strc.i[2]/sum(strc.i)*demrates$t12[i]
  gamma <- c(gamma, gamma.i)
  phi <- c(phi,phi.i)
  rho <- c(rho,rho.i)
}

# 4. calculate transition matrices for year 1:1001 (newborns are 1 years old)

surv <- NULL; fecu <- NULL
pb <- txtProgressBar(min=1,max=dim(demrates)[1],style=3)
for(i in 1:dim(demrates)[1]){
  matrix.i <- matrix(c(demrates$t11[i],demrates$t21[i],demrates$t12[i],demrates$t22[i]),2,2)
  F.i <- matrix(c(0,0,demrates$f12[i],0),2,2)
  matrix.ij <- diag(1,2)
  surv.i <- sum(matrix.ij[,1])
  fecu.i <- sum((F.i %*% matrix.ij/matrix(rep(apply(matrix.ij,2,sum),each=2),2,2))[,1])
  for(j in 1:1000){
```

```
    matrix.ij <- matrix.i %*% matrix.ij
    surv.i <- c(surv.i, sum(matrix.ij[,1]))
    fecu.ij <- (F.i %*% matrix.ij/matrix(rep(apply(matrix.ij,2,sum),each=2),2,2))[,1]
    fecu.ij[is.nan(fecu.ij)] <- 0
    fecu.i <- c(fecu.i, sum(fecu.ij))
  }
  surv <- rbind(surv,surv.i)
  fecu <- rbind(fecu,fecu.i)
  setTxtProgressBar(pb,i)
}

# 5. reproductive value

qx0 <- (t(apply((lx[,1000:1]*fecu[,1000:1]),1,cumsum)))[,1000:1] #age 1 to 1000

# 6. max age (Waples et al. 2013 Proc R Soc B)

max_age <- NULL
for(i in 1:dim(demrates)[1]){
  max_age.lx.i <- min(which(lx[i,]<0.01))-1
  max.age.qx.i <- min(which(qx0[i,]<0.01))-1
  max_age <- c(max_age,min(max_age.lx.i,max.age.qx.i))
}
summary(max_age);length(max_age)

# 7. calculate H, S, T

H <- NULL; S <- NULL; gentime <- NULL
pb <- txtProgressBar(min=1,max=dim(demrates)[1],style=3)
for(i in 1:dim(demrates)[1]){
  surv.i <- surv[i,]
  H.ii <- log(surv.i)*surv.i
  H.ii[is.na(H.ii)] <- 0
  H.i <- -cumsum(H.ii)/cumsum(surv.i)
  fecu.i <- fecu[i,]
  S.ii <- log(surv.i*fecu.i)*surv.i*fecu.i
  S.ii[is.na(S.ii)] <- 0
  S.i <- -cumsum(S.ii)
  T.i <- cumsum(surv.i*fecu.i*1:1001)/cumsum(surv.i*fecu.i)
  H <- c(H,H.i[max_age[i]])
  S <- c(S,S.i[max_age[i]])
  gentime <- c(gentime,T.i[max_age[i]])
  setTxtProgressBar(pb,i)
}

# 8. summary

lhtraits <- cbind(gentime,H,La,gamma,rho,phi,S,Lw)
lhtraits <- as.data.frame(lhtraits)
names(lhtraits) <- c("gentime","H","La","gamma","rho","phi","S","Lw")
write.csv(lhtraits,"life_history_traits.csv",row.names=F)


##------------------------------------
## 5. PCA of life history traits
##------------------------------------

# 1. PCA
lhtraits.pca <- prcomp(lhtraits, scale=T)
write.csv(lhtraits.pca$x,"life_history_traits_PCs.csv",row.names=F)
write.csv(lhtraits.pca$rotation,"life_history_traits_rotation.csv",row.names=T)

# 2. Variance explained
plot(lhtraits.pca)
summary(lhtraits.pca)
variance.pca <- apply(lhtraits.pca$x,2,var)

par(lwd=3,mgp=c(3,1.5,0),mar=c(5,5,1,1))
plot(1:8,variance.pca,cex=3,type="b",
     xaxt="n",cex.axis=2,las=1,xlab="",ylab="")
axis(1,1:8,paste("PC",1:8,sep=""),cex.axis=2)
par(lwd=1,mgp=c(3,1,0),mar=c(4.1,5.1,5.1,2.1))

# 3. Biplot
biplot(lhtraits.pca)
summary(lhtraits.pca$x[,1:2])

par(lwd=3,mar=rep(5,4),mgp=c(3,1.5,0))
plot(lhtraits.pca$x[,1],lhtraits.pca$x[,2],bg=cols.eta,cex=3,pch=21,lwd=2,col="gray50",
     cex.axis=2,cex.lab=2,las=1,xlab="",ylab="",yaxt="n",xaxt="n",
     xlim=c(-6,5),ylim=c(-4,4))
axis(1,c(-2:1*5),c(-2:1*5),cex.axis=2,las=1, lwd=3)
axis(2,c(-3,0,3),c(-3,0,3),cex.axis=2,las=1, lwd=3)
arrows(0,0,lhtraits.pca$rotation[,1]*6,lhtraits.pca$rotation[,2]*4,lwd=6,length=0.2,angle=20,col="black")
axis(3,c(-2:2*0.5*6),c(-2:2*0.5),cex.axis=2,las=1, lwd=3)
axis(4,c(-2:2*0.5*4),round(c(-2:2*0.5),1),cex.axis=2,las=1, lwd=3)
par(lwd=1,mar=c(4.1,5.1,5.1,2.1),mgp=c(3,1,0))


##------------------------------------
## 6. Elasticity analysis
##------------------------------------

library(popbio)
library(Ternary)

# 1. Elasticity
```

```
elast <- NULL
for(i in 1:dim(demrates)[1]){
  elast.i <- c(as.vector(sensitivity(matrix(as.numeric(demrates[i,c(1,2,6,3)]),2,2))*
                           matrix(as.numeric(demrates[i,c(1,2,4,3)]),2,2)),
               as.vector(sensitivity(matrix(as.numeric(demrates[i,c(1,2,6,3)]),2,2))*
                           matrix(as.numeric(demrates[i,c(1,2,5,3)]),2,2))[3])
  elast <- rbind(elast,elast.i)
}

colnames(elast) <- c("t11","t21","t12","t22","f12")
rownames(elast) <- NULL
elast <- as.data.frame(elast)
elast$growth <- elast$t21
elast$stasis <- elast$t11+elast$t12+elast$t22
elast$reproduction <- elast$f12

# 2. Ternary diagram

par(mar=rep(3,4))
TernaryPlot(point = 'up', alab = 'Growth', blab = 'Stasis', clab = 'Reproduction',
            grid.lines = 2, grid.lty = 3, grid.lwd = 3, grid.minor.lines = 4,
            grid.minor.lty = 3, grid.minor.lwd = 3,
            axis.cex=2, axis.lwd=3, axis.labels=seq(0,1,by=0.5), lab.cex=2, lab.offset = 0.16)
AddToTernary(points, elast[,6:8], bg=cols.eta, col="gray50", cex=2.5, pch=21, lwd=2)
par(mar=c(4.1,5.1,5.1,2.1))
```
